# What Is a Generative Model? Definitions, Disagreements, and Evaluation in Human Neuroimaging

**DOI:** 10.64898/2026.01.13.698266

**Authors:** Matthew D. Greaves, Leonardo Novelli, Michael Breakspear, Adeel Razi

## Abstract

The term *generative model* is widely used in human neuroimaging; however, its meaning is often left implicit. Prompted by observations and discussions at the 2025 Organization for Human Brain Mapping (OHBM) Annual Meeting, we surveyed members of the neuroimaging community to examine how generative models are defined, used, and evaluated in practice. Responses revealed some agreement on functional criteria— such as a model’s ability to simulate data—alongside marked disagreement about whether specific, widely used methods should be considered generative models. Evaluative priorities also varied across respondents, though out-of-sample generalization and interpretability were consistently emphasized. Rather than proposing a single definition, this perspective highlights the diversity of current usage and argues for greater clarity when the term is invoked.

## Main

To the authors of this perspective piece, the 2025 Organization for Human Brain Mapping (OHBM) Annual Meeting was notable for the ease with which the term *generative model* circulated—often appearing without explanation, and as though its meaning were self-evident. In the educational course on neural field theory, generative models appeared as mechanistic, biophysical descriptions of neuronal dynamics. In keynote talks and symposia, the same term was used to describe (generic) general linear models (GLMs), and black-box artificial intelligence systems. Elsewhere, in a roundtable celebrating the thirtieth anniversary of the statistical parametric mapping (SPM) toolbox, “generative modeling” was described as the defining feature that distinguished SPM from other neuroimaging software^*^.

What caught our attention was not just the breadth of these uses, but the tension between them. Generative models seemed, at once, to be something very general—almost a synonym for “model”—and something highly specific, even privileged. They were common, yet they were also invoked as a mark of conceptual or methodological distinction. This apparent contradiction led us to ask colleagues, informally (over coffee), what they understood the term to mean. Several of these conversations ended with the same suggestion: that we should survey the community in a more formal manner^†^.

It is worth briefly noting that other researchers have remarked on the ambiguity surrounding this term^2^, and that—without attempting a sociolinguistic analysis—this ambiguity appears to reflect a gradual drift in usage across disciplines. In machine learning, the term was associated with early work on Boltzmann machines and related probabilistic models^3,4^, though it appears earlier still in linguistics, where it referred to rule-based formalisms for producing well-formed linguistic structures^5^. In Bayesian statistics, the term came to denote the specification of a joint probability distribution over observed data and latent variables, with priors encoding assumptions about the data-generating process^6,7^. In parallel, influential work in computational neuroscience framed perception itself as a process of inference over latent causes of sensory input^8^. Subsequently, the term became associated with mechanistic and biophysical models—models in which parameters are intended to correspond, at least approximately, to known biological processes that give rise to measured signals—regardless of whether those models are cast in explicitly Bayesian terms^9,10^. More recently, with the increased visibility and use of large language models, the same term may now be more often understood to refer to artificial intelligence systems that produce text and other content^11^.

This is all to say that, from certain perspectives, a model may be considered generative not because it generates realistic synthetic data, but because it supports inference about latent causes or parameters assumed to have generated certain features of observed data. In neuroimaging, this can mean specifying how unobserved neuronal states, network interactions, or coupling weights give rise to second-order features of neural signals (correlations, for example) well enough to estimate those quantities, with realistic time-domain simulation not serving as the primary basis for evaluation^12–14^. While a diversity of perspectives is neither surprising nor undesirable, it does raise the possibility that researchers may be talking past one another—using the same words to refer to models that differ substantially in their aims, assumptions, and standards of evaluation. To move beyond anecdote and to characterize how these different understandings are distributed in practice, we conducted a short survey of the OHBM community, combining open-ended and structured questions about how generative models are defined, used, and evaluated. In what follows, we summarize the results of this survey and reflect on what they reveal about shared intuitions and points of divergence within the field.

## Results

Our survey comprised 14 questions in total, including a mixture of free-text responses, multiple-choice items, and rank-ordering questions (see Supporting Information). The survey was circulated via the OHBM mailing list and remained open for 2 weeks, with participation entirely voluntary; respondents were free to skip any question. For the exploratory, descriptive analyses reported here, we focused on respondents who completed at least 5% of the survey, yielding a final sample of 88 respondents; full methodological details are provided in the Supporting Information. Of those respondents who provided demographic information, 44.74% identified as principal investigators or group leaders, 28.95% as postdoctoral researchers, and 21.05% as doctoral students, with smaller proportions reporting roles such as research staff, clinicians, or other positions. Reported ages ranged from 25–34 years to 65 years or older, with the largest proportion of respondents (40.79%) falling within the 25–34-year age group. Among respondents who reported gender, 59.21% identified as men and 32.90% as women, with 7.89% preferring not to disclose. Respondents who provided location information were based across 15 countries, most commonly the United States (29.23%) and Australia (26.15%), followed by the United Kingdom (9.23%), Germany (6.15%), and Italy (6.15%), with additional representation from Canada, South Korea, India, Belgium, Switzerland, France, Finland, and Singapore.

In terms of primary research area, respondents most frequently reported working in magnetic resonance imaging (MRI) (43.59%) and methods development or computational modeling (21.79%), followed by multimodal imaging (12.82%) and electroencephalography (8.97%), with smaller representations from clinical and translational neuroscience, positron emission tomography, and magnetoencephalography. When asked how generative models are used in their own work (multiple selections permitted), respondents most frequently reported applications in mechanistic or biophysical modeling (48.72%) and whole-brain dynamical modeling (42.31%). Substantial proportions also reported using generative models for synthetic data generation or augmentation (37.18%) and clinical prediction or stratification (34.62%). Less frequently reported uses included encoding or decoding models (30.77%), normative modeling or deviation mapping (24.36%), and simulation-based or likelihood-free inference (10.26%). Taken together, these characteristics suggest that the survey captured a reasonably broad snapshot of the OHBM community.

### How do researchers define a “generative model”?

We conducted a thematic analysis of responses to the open-ended question “In one or two sentences: what is a generative model, in your own words?”, drawing on established qualitative methods^15^, in which all free-text responses were inductively coded, codes were iteratively refined, and higher-level themes were identified through grouping of related codes (see Supporting Information). Responses clustered into five themes, defining generative models as: (i) mechanistic or biophysical forward models, in which latent physiological states are mapped to observed data; (ii) probabilistic or Bayesian latent-variable models, specified via joint distributions over hidden causes and observations; (iii) models defined by their capacity to simulate data, irrespective of underlying mechanism; (iv) content-producing artificial intelligence systems, such as large language or image-generation models; and (v) hybrid or mixed definitions combining elements of multiple perspectives. This diversity was also reflected in responses to the multiple-choice question “Which definition best matches your use of generative model?” (Fig. 1a). The most commonly endorsed definition—”any model that can produce synthetic data, by any method (mechanistic, probabilistic, or machine learning based)”— placed particular weight on a model’s capacity to simulate data. In line with this emphasis, most respondents (66.28%) indicated that an explicit likelihood was *not* required for a model to count as generative, and most respondents (59.30%) indicated that models unable to generate raw data—operating only at the level of summary statistics—would *not* be considered generative.

**Figure 1.**
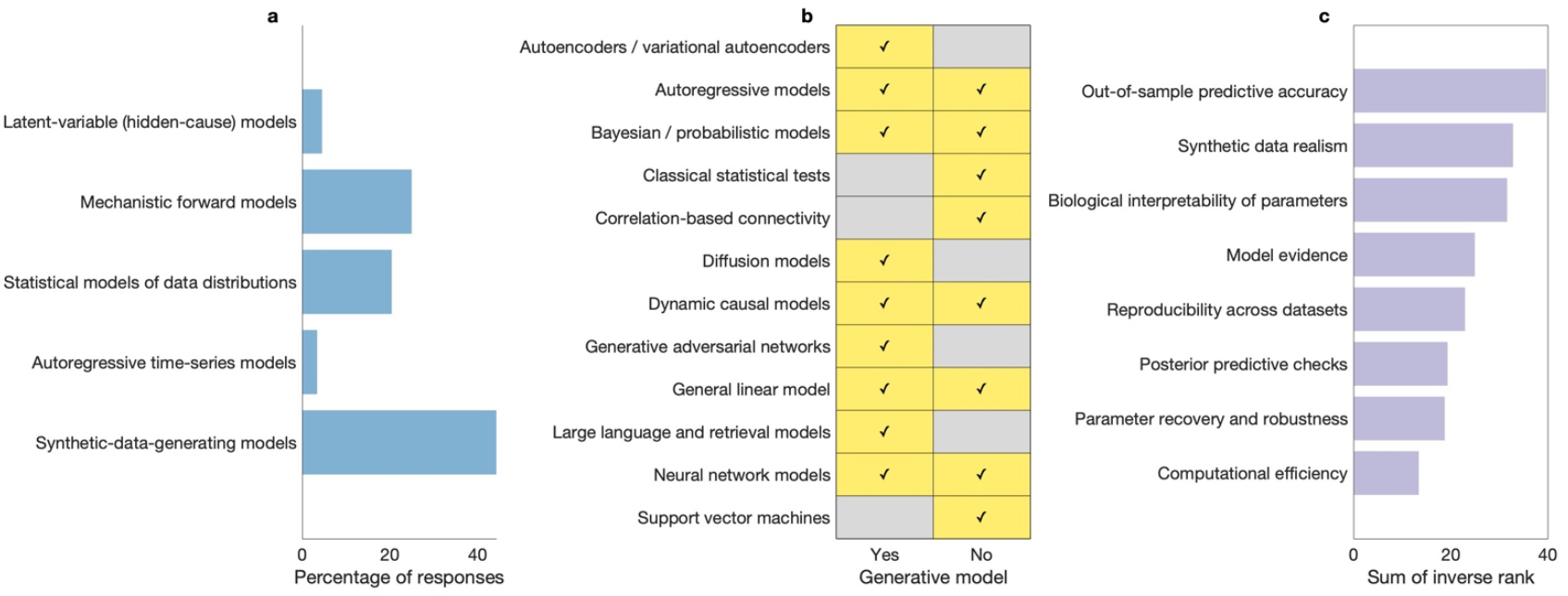
Definitions and evaluative criteria for generative models in neuroimaging. (**a**) Distribution of responses to the single-select multiple-choice question “Which definition best matches your use of generative model?”. The most frequently endorsed definition emphasized a model’s capacity to generate synthetic data, irrespective of mechanism. Percentages indicate the proportion of respondents endorsing each category. Two “Other” responses (not shown) reflected non-use or uncertainty about model classification. (**b**) Models named by respondents as generative or not generative in response to an open-ended prompt. Closely related methods were collapsed into broader model classes for visualization (for example, different forms of neural network models or probabilistic approaches). A checkmark indicates that at least one respondent classified the model class as generative or not generative, respectively (checkmarks, or absence thereof, do not imply consensus). Model classes appearing in both columns reflect disagreement. (**c**) Aggregated rankings of what respondents consider evidence of a “good” generative model. Bars show the sum of inverse ranks across respondents, such that higher values indicate criteria more frequently ranked as most important. Respondents placed greatest weight on out-of-sample predictive accuracy and generalization, followed by the ability to generate realistic synthetic data and the interpretability of model parameters. Full question wording is provided in the Supplementary Information.

Despite this seeming convergence and liberality, responses to a question asking participants to name one model they considered generative and one they considered *not* generative revealed substantial disagreement. Many of the same approaches were cited on both sides of the generative/non-generative divide by different respondents (Fig. 1b). For example, dynamic causal models, autoregressive models, and the GLM were each described as generative by some participants and explicitly non-generative by others. These disagreements were typically justified by reference to different evaluative criteria, including whether a model specifies an explicit forward process, whether it can generate raw data, and whether its parameters are interpreted as causal or mechanistic. In contrast, contemporary machine-learning systems such as generative adversarial networks and large language models were more consistently described as generative, while purely discriminative or descriptive models were more often cited as non-generative.

### What do researchers consider evidence of a “good” generative model?

We conducted a thematic analysis of responses to the open-ended question “What is one best practice you would recommend to others when evaluating generative models?”. Responses clustered into six themes, emphasizing: (i) generalization and out-of-sample validation, including the use of independent datasets and safeguards against data leakage; (ii) interpretability and biological plausibility, particularly the meaningfulness and identifiability of model parameters; (iii) simulation realism and failure modes, including stress-testing models and examining when they break down; (iv) model comparison and falsifiability, through the use of alternative models, null models, or formal comparison criteria; (v) appropriateness of assumptions, including parsimony and alignment between model, data, and research question; and (vi) explicit resistance to codified best practices, reflecting concerns that prescriptive standards may constrain innovation or obscure context-specific goals. When asked to rank evidence for a “good” generative model, respondents placed the greatest weight on out-of-sample predictive accuracy and generalization, followed by a model’s ability to generate realistic synthetic data and the biological interpretability of its parameters (Fig. 1c).

### What do researchers consider to be misused terminology?

To assess whether ambiguity surrounding generative models reflects a broader issue of terminology in neuroimaging, we examined responses to the question asking participants to identify commonly misused modeling terms and to clarify how they would define them. Responses highlighted concerns about the use of language that implies mechanistic or biological realism without sufficient justification, the tendency to treat formal statistical criteria as guarantees of scientific validity, and the conflation of prediction, explanation, and interpretation. Several respondents also expressed unease with uncritical adoption of machine-learning terminology in neuroimaging contexts.

## Discussion

The aim of this perspective was deliberately modest. We set out to document how generative models are currently understood, used, and evaluated within the human neuroimaging community. Prompted by informal conversations at the 2025 OHBM Annual Meeting, we conducted a short survey combining open-ended and structured questions. Our key finding is that, while definitions of what constitutes a generative model varied widely across respondents, judgments about how such models should be evaluated tended to converge. While other authors have remarked on disagreements surrounding modeling concepts and terminology^16–20^, to our knowledge this is the first attempt to survey how a single, widely used term is interpreted within neuroimaging.

Several patterns emerge clearly. On the surface, respondents appeared to agree on broad criteria for what counts as a generative model—most commonly, the capacity to simulate data (Fig. 1a). Yet agreement was less apparent when participants were asked to classify specific methods: the same models were repeatedly placed on opposite sides of the generative–non-generative divide (Fig. 1b). In some cases, these disagreements appeared to reflect differences in context of use. A GLM, for example, might be considered a mechanistic model when used to map known experimental inputs—convolved with a hemodynamic response function—onto functional MRI data^21,22^, whereas in other settings it may simply be used as an ordinary least-squares tool for relating observed variables, without much commitment to how those relationships should be understood^23^. Despite divergent definitions, when it came to evaluative criteria respondents largely converged on the importance of out-of-sample generalization, simulation realism, and interpretability, while placing less weight on formal criteria such as model evidence or computational efficiency (Fig. 1c). Agreement, where it existed, tended to concern *what models should do*, rather than *what they should be called*.

The final set of responses helps to place these findings in a broader context. When asked about commonly misused terminology, participants pointed to similar problems elsewhere in the field: language that implies mechanistic or biological realism without sufficient justification, and persistent confusion between prediction, explanation, and interpretation (for example). If there is one practical recommendation to draw from this survey, it is simply this: when using the term *generative model* (and indeed, any other ambiguous term), researchers should state explicitly what they mean by it in context—whether they are referring to a probabilistic data-generating process, a mechanistic forward model, or something else. This will not enforce a single definition, but it can make explicit the assumptions and evaluative criteria underlying different uses of the term, with practical consequences for how modeling contributions are assessed.

Finally, it is worth noting that the disagreements captured by the survey were not entirely dispassionate. The tone of many free-text responses was striking. Some respondents expressed strong convictions about correct usage, often treating their position as self-evident. Others resisted the premise of the questions themselves, rejecting the idea that such distinctions require formal definition or best-practice guidelines. Alongside this, several participants openly described confusion. Taken together, these reactions suggest that ambiguity around generative models is not merely a technical problem, but one bound up with disciplinary identity, pedagogical norms, and assumptions about what should go without saying. Our aim here is not to settle these questions, but to make them visible. If this work contributes to clearer communication—especially for early-career researchers and interdisciplinary collaborators—then it will have served its purpose. Any future attempt to articulate shared guidelines will, inevitably, meet both agreement and resistance, and it should.

## Data and code availability statement

The survey data analyzed in this study have been de-identified and include only revised codes and thematic variables, with no free-text responses, initial coding labels, or direct identifiers retained. The processed dataset and all analysis code are publicly available at https://github.com/mdgreaves/ohbm-generative-models-survey, and are archived at Zenodo: https://doi.org/10.5281/zenodo.18238047, enabling full reproducibility of the reported results.

## Footnotes

* These observations are necessarily impressionistic, as not all talks were archived. However, all accepted abstracts from OHBM 2025 were deposited on Zenodo^1^, and a term search of this corpus returns multiple instances of “generative model” and related phrases, consistent with the breadth of usage described here.

† We are particularly grateful to Joseph Lizier for encouraging us to formalize these informal discussions and to survey the broader community.

## References

1. Aperture Neuro. OHBM 2025 Annual Meeting Abstract Book. Zenodo. Preprint posted online June 13, 2025. doi:10.5281/ZENODO.15641972

2. Maran R, Müller EJ, Fulcher BD. Analyzing the brain’s dynamic response to targeted stimulation using generative modeling. Network Neuroscience. 2025;9(1):237–258. doi:10.1162/netn_a_00433

3. Ackley D, Hinton G, Sejnowski T. A learning algorithm for boltzmann machines. Cognitive Science. 1985;9(1):147–169. doi:10.1016/S0364-0213(85)80012-4

4. Dayan P, Hinton GE, Neal RM, Zemel RS. The Helmholtz Machine. Neural Computation. 1995;7(5):889–904. doi:10.1162/neco.1995.7.5.889

5. Šaumjan SK. Outline of the applicational generative model for the description of language. Foundations of Language. Published online 1965:189–222.

6. Bishop CM. Pattern Recognition and Machine Learning. Springer; 2006.

7. Gelman A, Carlin JB, Stern HS, Dunson DB, Vehtari A, Rubin DB. Bayesian Data Analysis. 0 ed. Chapman and Hall/CRC; 2013. doi:10.1201/b16018

8. Rao RPN, Ballard DH. Predictive coding in the visual cortex: a functional interpretation of some extra-classical receptive-field effects. Nat Neurosci. 1999;2(1):79–87. doi:10.1038/4580

9. Breakspear M, Heitmann S, Daffertshofer A. Generative Models of Cortical Oscillations: Neurobiological Implications of the Kuramoto Model. Front Hum Neurosci. 2010;4. doi:10.3389/fnhum.2010.00190

10. Friston KJ, Harrison L, Penny W. Dynamic causal modelling. NeuroImage. 2003;19(4):1273–1302. doi:10.1016/S1053-8119(03)00202-7

11. Sengar SS, Hasan AB, Kumar S, Carroll F. Generative artificial intelligence: a systematic review and applications. Multimed Tools Appl. 2024;84(21):23661–23700. doi:10.1007/s11042-024-20016-1

12. Friston KJ, Kahan J, Biswal B, Razi A. A DCM for resting state fMRI. NeuroImage. 2014;94:396–407. doi:10.1016/j.neuroimage.2013.12.009

13. Frässle S, Lomakina EI, Razi A, Friston KJ, Buhmann JM, Stephan KE. Regression DCM for fMRI. NeuroImage. 2017;155:406–421. doi:10.1016/j.neuroimage.2017.02.090

14. Raj A, Sipes BS, Verma P, Mathalon DH, Biswal B, Nagarajan S. Spectral graph model for fMRI: A biophysical, connectivity-based generative model for the analysis of frequency-resolved resting-state fMRI. Imaging Neuroscience. 2024;2:imag-2-00381. doi:10.1162/imag_a_00381

15. Braun V, Clarke V. Using thematic analysis in psychology. Qualitative Research in Psychology. 2006;3(2):77–101. doi:10.1191/1478088706qp063oa

16. Levenstein D, Alvarez VA, Amarasingham A, et al. On the Role of Theory and Modeling in Neuroscience. J Neurosci. 2023;43(7):1074–1088. doi:10.1523/JNEUROSCI.1179-22.2022

17. Friston KJ. Functional and Effective Connectivity: A Review. Brain Connectivity. 2011;1(1):13–36. doi:10.1089/brain.2011.0008

18. Bzdok D, Ioannidis JPA. Exploration, Inference, and Prediction in Neuroscience and Biomedicine. Trends in Neurosciences. 2019;42(4):251–262. doi:10.1016/j.tins.2019.02.001

19. Razi A, Friston KJ. The Connected Brain: Causality, models, and intrinsic dynamics. IEEE Signal Process Mag. 2016;33(3):14–35. doi:10.1109/MSP.2015.2482121

20. Greaves MD, Novelli L, Mansour L. S, Zalesky A, Razi A. Structurally informed models of directed brain connectivity. Nat Rev Neurosci. 2025;26(1):23–41. doi:10.1038/s41583-024-00881-3

21. Friston KJ, Holmes AP, Worsley KJ, Poline J.P., Frith CD, Frackowiak RSJ. Statistical parametric maps in functional imaging: A general linear approach. Human Brain Mapping. 1994;2(4):189–210. doi:10.1002/hbm.460020402

22. Lindquist MA, Meng Loh J, Atlas LY, Wager TD. Modeling the hemodynamic response function in fMRI: Efficiency, bias and mis-modeling. NeuroImage. 2009;45(1):S187–S198. doi:10.1016/j.neuroimage.2008.10.065

23. Mumford JA, Nichols T. Simple group fMRI modeling and inference. NeuroImage. 2009;47(4):1469–1475. doi:10.1016/j.neuroimage.2009.05.034

