## Supporting Information for "What Is a Generative Model? Definitions, Disagreements, and Evaluation in Human Neuroimaging"


### Methods
#### Survey design and recruitment
The study comprised an online survey of the neuroimaging community designed to probe how generative models are defined, used, and evaluated in practice. The survey consisted of 14 questions in total, including a mixture of free-text responses, multiple-choice items, multi-select questions, and rank-ordering tasks. A full list of survey questions and response options is provided in the next section (see Survey). The survey was circulated via the Organization for Human Brain Mapping (OHBM) electronic mailing list and remained open for a period of two weeks. Participation was entirely voluntary, and respondents were free to skip any question. No incentives were offered for participation.

#### Ethics statement
The survey preamble informed participants of the study's aims, the voluntary nature of participation, and the intended use of responses in aggregate form. No incentives were offered. No personally identifying information was collected as part of the survey, except where respondents may have chosen to include such information in free-text responses. Formal ethics committee approval was not sought.

#### Sample inclusion
For the exploratory, descriptive analyses reported in the paper, we included respondents who completed at least 5% of the survey, yielding a final sample of 88 respondents. Because respondents were permitted to skip individual questions, sample sizes vary across questions. All such details are recoverable via the open-source analysis code.

#### Qualitative analysis of free-text responses
Free-text responses were collected in response to questions 1, 5 and 9 (see Survey). We conducted a thematic analysis following established qualitative methods[1], using an inductive coding approach. All free-text responses were initially coded without predefined categories. Codes were iteratively refined across the full set of responses to improve internal consistency and reduce redundancy. Higher-level themes were subsequently identified by grouping conceptually related codes. To reduce identifiability risks, initial fine-grained codes and raw free-text responses are not publicly shared. Instead, revised codes and thematic labels were generated and used for all reported analyses.

**Quantitative analysis**
Quantitative analyses focused on descriptive summaries of survey responses. Categorical variables (e.g., career stage, research area, country) were summarized using counts and percentages relative to the number of respondents for each item. For multi-select questions, responses were parsed by matching canonical response options within each respondent's selection string. Percentages were computed as the proportion of respondents selecting each option (rather than the proportion of total selections). For rank-ordering questions, responses were aggregated using an inverse-rank scoring scheme, in which lower numerical ranks (indicating higher preference) contributed larger weights. Specifically, for each criterion, inverse ranks were summed across respondents, yielding an aggregate score used solely for relative ordering and visualization. No inferential statistical tests were performed; all analyses are descriptive in nature.

**Data and code availability**
The survey data analyzed in this study have been de-identified and include only revised codes and thematic variables, with no free-text responses, initial coding labels, or direct identifiers retained. The processed dataset and all analysis code are publicly available at https://github.com/mdgreaves/ohbm-generative-models-survey, and are archived at Zenodo: https://doi.org/10.5281/zenodo.18238047, enabling full reproducibility of the reported results.

### Survey

Preamble:
Welcome to the Generative Models in Neuroimaging Survey

Thank you for your interest in participating.

This survey aims to assess how researchers across neuroimaging and computational neuroscience define, use, and evaluate generative models. The term 'generative model' is applied in many different ways—to small-scale biophysical models and large-scale machine-learning-based approaches—and we hope to capture this diversity of perspectives.

Your responses will help us map current practices, clarify terminology, and identify areas where methodological guidelines may be needed.

Estimated time to complete: 5–10 minutes
(The survey contains a mixture of multiple-choice and short free-text responses.)

Participation is voluntary, and you may skip any question. No identifiable personal information is collected unless you choose to provide it in a free-text response. Responses will be analysed in aggregate for a publication on generative modelling in neuroimaging.

If you have questions, please feel free to contact Adeel Razi:

Thank you for contributing your expertise.

Questions:
1. In one or two sentences: What is a "generative model" in your own words? (Free text)
2. Which definition best matches your use of "generative model"? (Multiple-choice; single select)
    a. A mechanistic model that simulates observed data via an explicit forward process (e.g., neuronal–haemodynamic model, biophysical state-space model, dynamic causal modelling).
    b. A statistical model that learns a data distribution and can sample new data (e.g., variational autoencoder, diffusion models, generative adversarial networks).

c. A latent-variable model that explains data in terms of hidden causes regardless of whether it explicitly simulates data (e.g., Bayesian causal model, hierarchical latent-cause models).

d. A statistical time-series model that generates data via dependencies on past values (e.g., autoregressive model, autoregressive moving average model).

e. Any model that can produce synthetic data, by any method (mechanistic, probabilistic, or machine learning based).

f. Other (please specify).

3. Does a model need an explicit likelihood (i.e., a specified $p(x/\theta)$) to count as generative? (Yes/No)
4. If a model cannot simulate raw data (only summary statistics), do you still consider it generative? (Yes/No)
5. Name one method you consider generative and one you do not, and briefly explain why. (Free text)
6. Give an example of terminology that you think is commonly misused in the context of modeling and neuroimaging, and how you would define it. (Free text)
7. Which of the following best describes the use of generative models in your work? (Checkboxes; multi-select)

   a. Mechanistic/biophysical modelling (e.g., neuronal–haemodynamic models, state-space models, dynamic causal models)

   b. Whole-brain dynamical modelling (e.g., network-level simulations, eigenmode-based models)

   c. Synthetic data/augmentation (e.g., variational autoencoders, generative adversarial networks, diffusion models, simulation pipelines)

   d. Normative modelling/deviation maps (e.g., hierarchical Bayesian normative models, Gaussian processes)

   e. Encoding/decoding models (e.g., generative encoding models, latent-variable models)

   f. Clinical prediction/stratification (e.g., individualised latent-space models, disease subtyping)

   g. Simulation-based inference / likelihood-free inference

   h. Other (please specify)
8. Please rank the following as evidence of a ‘good’ generative model in your area of neuroimaging: (Rank order)

   a. Out-of-sample predictive accuracy / generalisation performance

   b. Model evidence (e.g., marginal likelihood, variational free energy)

   c. Posterior predictive checks / simulation-based calibration

   d. Parameter recovery / identifiability / robustness to noise

   e. Ability to generate realistic synthetic data

   f. Biological interpretability and meaningfulness of parameters

   g. Computational efficiency / tractability

   h. Stability across datasets / reproducibility
9. What is one best practice you would recommend to others when evaluating generative models? (Free text)

The following questions are to help us describe who took this survey.
All questions are optional.

1. Primary research area: (Single select)

   a. MRI

   b. EEG

   c. MEG

   d. PET

   e. fNIRS

   f. Multimodal imaging

   g. Methods development / computational modelling

   h. Clinical research / translation neuroscience

   i. Other (please specify)

2. Primary role: (Single select)
    a. PI / Group leader
    b. Postdoctoral researcher
    c. PhD Student
    d. Masters Student
    e. Research Assistant / Technician / Data Scientist
    f. Clinician (e.g., psychiatrist, neurologist, radiologist)
    g. Other (please specify)
3. Which age group are you in? (Single select)
    a. Under 18
    b. 18–24
    c. 25–34
    d. 35–44
    e. 45–54
    f. 55–64
    g. 65 or older
    h. Prefer not to say
4. Which of the following best describes your gender? (Single select)
    a. Woman
    b. Man
    c. Non-binary
    d. Prefer not to say
    e. Other (please specify)
5. In which country do you currently work or study? (Single select)

### References


1. Braun V, Clarke V. Using thematic analysis in psychology. *Qualitative Research in Psychology*. 2006;3(2):77-101. doi:10.1191/1478088706qp063oa
